# Automatic semantic segmentation of the osseous structures of the paranasal sinuses

**DOI:** 10.1101/2024.06.21.599833

**Authors:** Yichun Sun, Alejandro Guerrero-López, Julián D. Arias-Londoño, Juan I. Godino-Llorente

## Abstract

Endoscopic sinus and skull base surgeries require the use of precise neuronavigation techniques, which may take advantage of accurate delimitation of surrounding structures. This delimitation is critical for robotic-assisted surgery procedures to limit volumes of no resection. In this respect, accurate segmentation of the Osseous Structures surrounding the Paranasal Sinuses (OSPS) is a relevant issue to protect critical anatomic structures during these surgeries. Currently, manual segmentation of these structures is a labour-intensive task and requires expertise, often leading to inconsistencies. This is due to the lack of publicly available automatic models specifically tailored for the automatic delineation of the complex OSPS. To address this gap, we introduce an open-source data/model for the segmentation of these complex structures. The initial model was trained on nine complete ex vivo CT scans of the paranasal region and then improved with semi-supervised learning techniques. When tested on an external data set recorded under different conditions and with various scanners, it achieved a DICE score of 94.82 *±* 0.9. These results underscore the effectiveness of the model and its potential for broader research applications. By providing both the dataset and the model publicly available, this work aims to catalyse further research that could improve the precision of clinical interventions of endoscopic sinus and skull-based surgeries.

## I. INTRODUCTION

**P**ARANASAL sinus surgery, particularly as a treatment for rhinosinusitis or the removal of nasal polyps, has become frequent [1]. These surgeries use endoscopic techniques that are approached through the nasal cavity [2]. Alternatively, skull-based surgeries, such as pituitary tumours, meningiomas, or acoustic neuromas, are also procedures that are approached —in most cases— through the nasal cavity using endoscopic techniques [3]. Due to the proximity of the paranasal area to the ocular orbit and cranial nerves, precision in these surgeries is crucial to avoid major complications such as blindness, central nervous system injuries, trauma, and even death. This underscores the importance of high-precision navigation procedures to minimise risks and ensure optimal outcomes [4].

The support of computerised navigation systems to approach endoscopic sinus and skull base surgeries is a common approach to achieve the required level of precision [5]. Modern neuronavigation systems allow real-time monitoring of the position of the endoscopic instrument relative to the three planes of the preoperative Computed Tomography (CT) or Magnetic Resonance (MR) scans of the patient [6]. The information provided by the neuronavigation tools can also be complemented with acoustic or haptic feedback when the boundaries of a certain region of interest are reached [7].

CT imaging greatly improved the understanding of sinonasal anatomy and pathology since its introduction [8]. This high-resolution imaging technique is essential not only for identifying anatomical variants [9] but also for precise surgical planning in sinus and skull-based surgeries. Its application directly influences outcomes in treating conditions such as nasopharyngeal cancer [10], obstructive sleep apnea [11], and rhinosinusitis [12]. Additionally, CT imaging is key for planning surgeries for skull base tumours, including fossa meningioma [13], enhancing surgical safety.

Detailed segmentation of the osseous structures surrounding the skull base and paranasal sinuses (Fig. 1) can significantly improve the characteristics of current neuronavigators by establishing clear boundaries for volumes that cannot be resected, thus reducing surgical complications. However, the complex anatomical structures of the paranasal sinuses and their significant variability between individuals [9] pose particular challenges not yet solved.

**Fig. 1.**
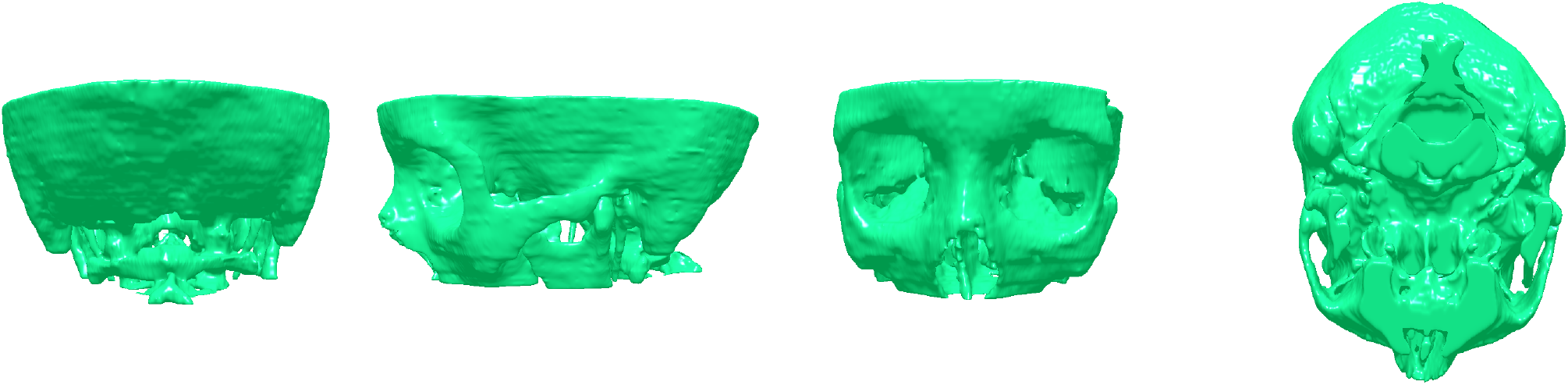
3D reconstruction of the OSPS. From left to right: back, lateral, front, and bottom views

Furthermore, delimiting anatomical structures is especially critical for robot-assisted endoscopic sinus and skull base surgeries, where precise identification of 3D boundaries is essential [14]. By accurately mapping osseous structures, surgeons can navigate complex anatomical volumes adjacent to the paranasal sinuses more effectively, thus minimising the risks associated with these invasive procedures [15]. This task is typically carried out manually or semi-automatically (i.e., automatically but supervised by human experts), which is considered the gold standard for this purpose. This is a laborious procedure due to the complexity and size of the tiny structures to be delineated, and the large number of slices available for each CT scan [16]. Besides, it requires high levels of expertise [17], is very time-consuming, and introduces a certain variability due to different delineation criteria introduced by different experts [18]. Consequently, new automatic methods are required to segment the aforementioned structures.

In recent years, driven by advances in Deep Learning (DL) and image processing, substantial progress has been made in the automatic segmentation of many body structures and/or tissues using CT scans [19]. These advances are significant in delineating osseous structures [20], which in some cases are relatively straightforward to segment due to their size, density, and contrast with respect to their surrounding tissues. In this respect, large bones, such as the femur, tibia or fibula, are typically simple to segment [21] achieving Sørensen-Dice coefficient (DICE) scores up to 97.28 *±* 1.73. Ribs present more complications [22], scoring with a DICE up to 94.9. Further complex targets include the segmentation of maxillofacial structures [23], [24], with DICE scores ranging from 82 to 94. In addition, studies in [25] and [26] present a much more challenging segmentation of the associated tiny structures in the inner ear, reporting scores that range from 56.0 for the stapes to 95.2 for the labyrinth.

### A. The automatic segmentation of the cranial structures of the paranasal sinuses. Related works

The works in [23] and [24] present automatic models for the segmentation of the upper skull, including the entire volume from the crown to the maxilla. The authors used advanced neural networks such as nnU-Net and modified U-Nets, achieving DICE scores of 96.2 and 96.5, respectively. These scores are biased by the better precision obtained for large skull bones, such as frontal, parietal, and occipital. Consequently, do not represent the specific precision achieved in the segmentation of the complex osseous paranasal structures.

In addition, [27] reports a DICE score of 78.8 *±* 10.3 using a 3D U-Net transformer-based (UNETR), for complex structures of the cranial base that include some of the adjacent regions of the paranasal sinuses. Furthermore, a DICE score of 94.0 for the skull base is reported in [28] using a 3D U-Net.

In the context of sinus scan analysis, existing research focusses mainly on segmenting the upper airway rather than osseous structures. In this regard, [29] achieved a DICE score of 90.7 for the maxillary sinus. Other approaches reported DICE scores up to 93.0 for the entire upper airway (including all paranasal sinuses) using a 3D U-Net [28], [30]. These results are of obvious interest, but the strong contrast of the upper airway significantly reduces the complexity of the challenge.

In summary, recent research on automatic segmentation of the osseous structures of the skull focusses on the large cranial bones. However, the specific challenges of delineating the OSPS (Fig. 1) are scarcely addressed. Thus, new automatic methods are required to segment these intricate structures. In addition, to our knowledge, there are no open datasets of CT scans specifically annotated with the OSPS. This is in part attributed to the specialised and hard work required to delineate each scan [16]. As an example, and to illustrate the effort required to manually annotate the sinus volume, the work in [31] reports an average of 13 hours for a semi-automatic segmentation of the upper airway. Due to the complexity of the intricate osseous structures of the paranasal sinuses, the time required to segment them is expected to be longer than that dedicated to the upper airway.

## II. MATERIAL AND METHODS

This section describes the material and methods, starting from the datasets, the annotation process to create the masks, the methods for pre-processing the images, the architectures used, and the experimental protocol.

### A. Datasets

This section introduces two corpora used for the Experimental Phase (EP). These datasets were retrospectively collected in 2012, and have been used since then in different research projects and with different objectives. Data collection was carried out according to strict ethical protocols. In the context of this paper, the two corpora are referred next as *internal* and *external* datasets.

All CT scans are axial views stored in Digital Imaging and Communications in Medicine (DICOM) format. The scanners are spiral and correspond to four different models from three manufacturers. Slices were reconstructed using six distinct kernels that were fixed during the acquisition process. All slices are 2D greyscale images, each with a resolution of 512×512 pixels. Table I summarises the most relevant characteristics of each dataset. Detailed information corresponding to each CT scan and patient can be accessed in the public repository associated with this work.

**TABLE I.**
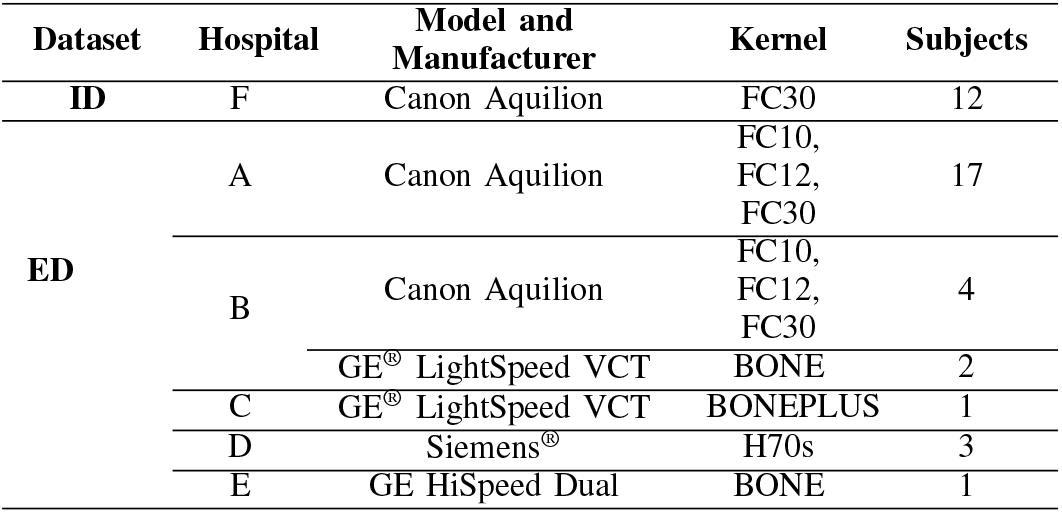
CHARACTERISTICS OF THE INTERNAL AND EXTERNAL DATASETS.

#### a) Internal Dataset (ID)

Comprises full head CT ex vivo scans obtained post mortem from twelve individuals who donated their organs for research purposes (see Fig. 2.Top). They were recorded using a Canon^®^ Aquilion scanner. Tissues were treated with specific preservative treatments, which sometimes partially fill the upper airway but do not affect the cranial structures in any sense. Each scan contains slices with a thickness of 1.00 mm. These scans cover the whole head volume, from the crown to the neck.

**Fig. 2.**
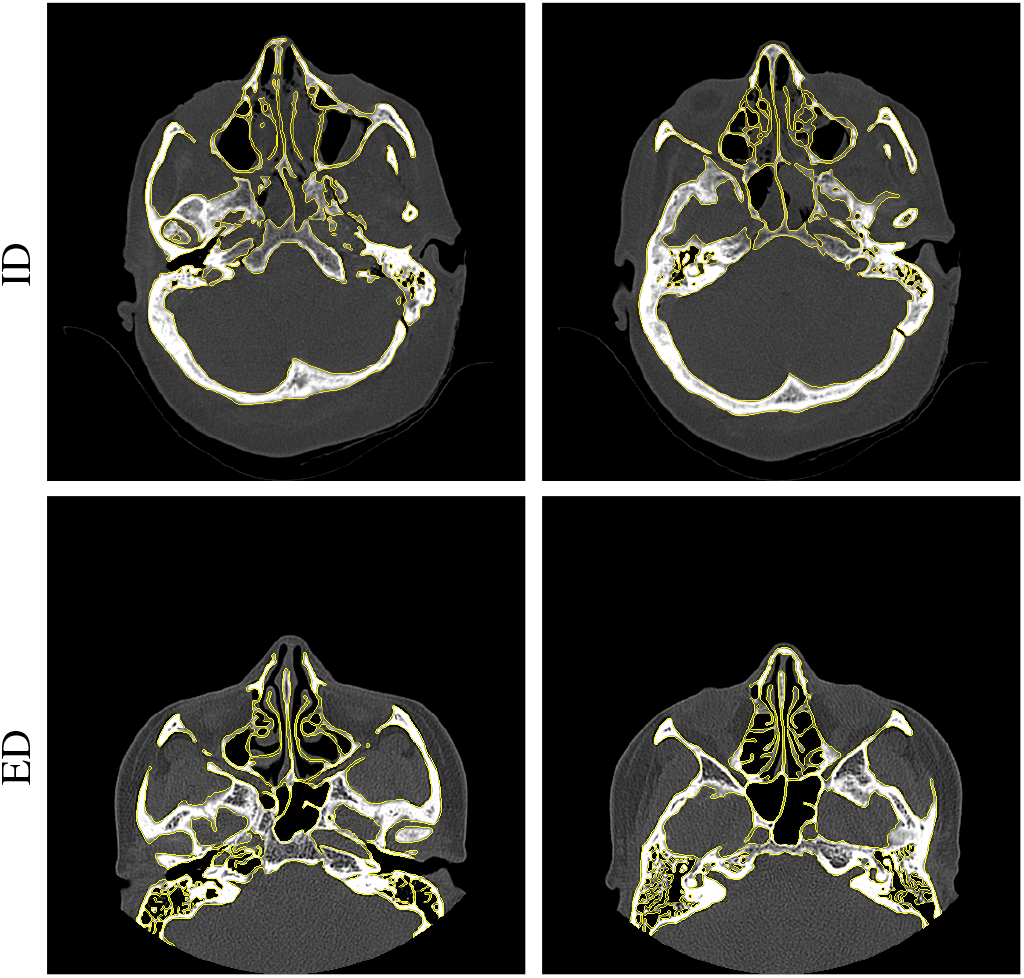
Examples of manual segmentation of the OSPS. Figures in the top correspond with slices taken from the *ID* (full head scans). Figures in the bottom correspond with slices taken from the *ED* (sinus scans). Note the differences in the field of *view*.

#### b) External Dataset (ED)

The second dataset includes in vivo CT sinus scans^1^ (i.e., with a field of view restricted to the sinus area (Fig. 2.Down), and with a range from the upper alveolar bone to the frontal sinuses), which were collected during clinical routine. They correspond to forty additional patients (with an average age of 29 *±* 16 yrs. old) who agreed on the use of their data for research purposes. The scans were obtained using four different scanners from four different hospitals (as detailed in Table I), and were retrieved from their corresponding Picture Archiving and Communication System (PACS). The slice thickness ranged from 0.47 to 1.00 mm, with a mean value of 0.59*±*0.17 mm.

### B. Data annotation

An annotation procedure is required to create the masks needed for the training and validation of the models. The gold standard is a manual delimitation. This is a labour-intensive step that requires extensive knowledge of the anatomical structures of the skull (see Section I-A). The structures of interest were manually delineated with 1-pixel width contours (Fig. 3), and the bowels were subsequently filled to obtain the final masks. Fig. 3 exemplifies the complexity of manual annotation. A different manual annotation procedure was used for each dataset:

**Fig. 3.**
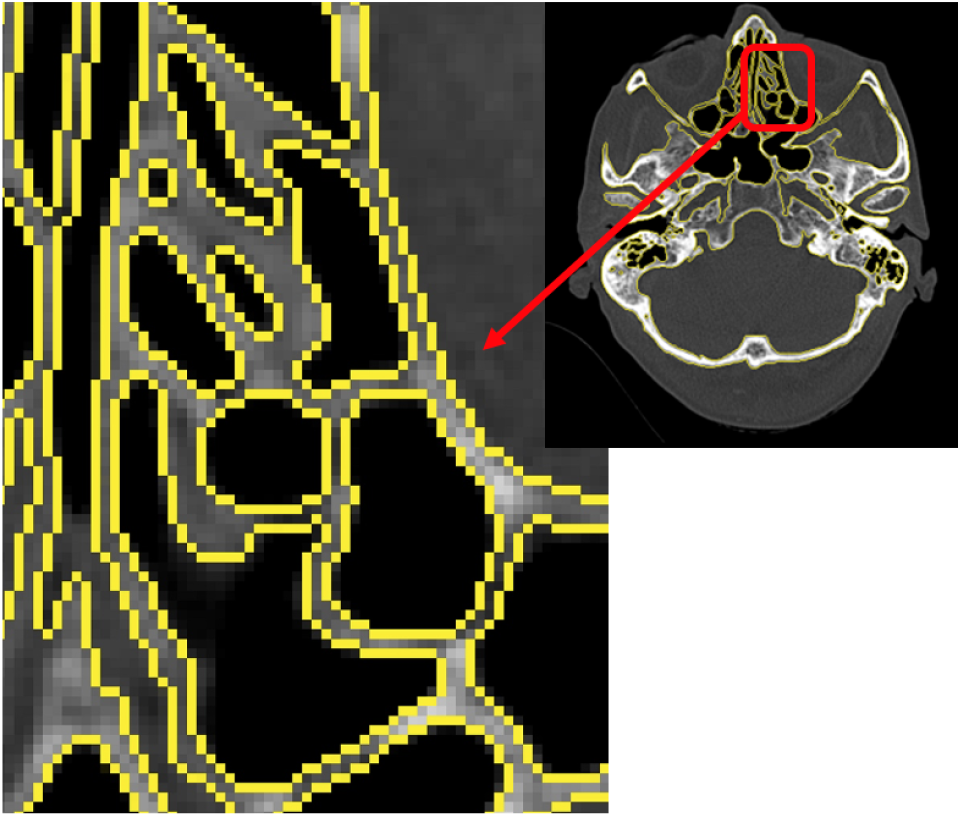
Detail of the contours manually delineated in an area corresponding to the ethmoidal sinuses. The left image is a zoomed-in view of the red box on the right. The structures are carefully delineated using 1-pixel width contours. Note that some of the structures to be segmented are also 1 or 2 pixels width.

#### a) ID

Nine out of the twelve scans available in the ID were manually segmented from scratch by two experts (i.e., no semi-automatic methods were used). One of the experts carried out a reviewing process of all manual contours to ensure the consistency of the masks created. Axial slices annotated are those in the volume of interest (i.e., from the upper alveolar bone to the frontal sinuses, including the entire paranasal sinuses region). This volume is covered by 64 adjacent slices. The slices of each of the nine subjects, along with their corresponding masks, are collectively designated as Subset 1 (S1). Due to the already commented complexity of the structures to be delineated, the annotation time ranged from 40 to 90 min. per slice (with an average time of 63 min.), i.e., 80 hours per volume.

#### b) ED

For the ED, manual masks were delineated by seven different experts. The expert who carried out the reviewing process of the ID reviewed the consistency and accuracy of the masks created in this scenario. A total of 4 CT scans out of 28 were partially annotated. These scans were chosen to correspond to the 4 different CT machines available in the dataset and 4 different hospitals, specifically, A, B, D and E.

Aligned with the state-of-the-art [31], [32], the annotation methods applied to the ED followed a semi-automatic procedure. The masks were extracted using the best-trained model with the ID (see Section II-H), and manually corrected pixel-wise. The number of slices selected depends on the thickness: 64 for those scans with a thickness of 1.00 mm., 85 when it was 0.75 mm., and 128 when it was 0.50 mm. Subsequently, within this range, 30 discontinuous slices were manually annotated. These are spaced with 1-4 unannotated slices, but the volume covered is still from the upper alveolar bone to the frontal sinuses. These manually annotated slices are collectively designated as Subset 2 (S2). The manual annotation took between 15 and 50 min. per slice (25 min. on average). The remaining slices in the ED —not included in S2—, have been automatically segmented by the model trained using S1. In contrast, the non-annotated slices of both datasets have been collectively designated as Subset 3 (S3). Table II provides detailed information about the annotated data and their distribution across the subsets.

**TABLE II.** DETAILS OF DATA ANNOTATION AND DISTRIBUTION OF SUBSETS.

| Dataset | Subjects | Subset | Slices/Patient | Manual masks/Patient |
| --- | --- | --- | --- | --- |
| **ID** | 9 | S1 | 64 | 64 continuous |
| **ED** | 4 | S2 | 64 | 30 discontinuous |
|  |  |  | 64 | 30 discontinuous |
|  |  |  | 85 | 30 discontinuous |
|  |  |  | 128 | 30 discontinuous |
| **ID & ED** | 27 | S3 | all | 0 |

In summary, S1, derived from the ID, contains a total of 576 slices and their corresponding manual masks. S2, from the ED, comprises 120 slices and their corresponding masks (excluding unannotated ones). Together, this results in a total of 696 samples available with ground truth.

### C. Data preprocessing

All DICOM files were converted to 2D PNG images (one for each slice). In addition, all CT scans were standardised with the same window size (represented by its Window Level (WL) and Window Width (WW)). The window size was initially fixed as [WL, WW]=[500, 2000] for all CT scans in both datasets. Subsequently, images were normalised and equalised. Greyscale images of one channel were scaled in the range [0,1]. Then, Contrast Limited Adaptive Histogram Equalisation (CLAHE) was used to enhance contrast [33]– [35]. The image was divided into 8 *×* 8 small blocks, each of which was equalised independently. In this respect, the contrast limit threshold was set to 2.0. Furthermore, the slices of each subject were converted to a 3D volume after stacking (3DVAS) for training 3D models. This preprocessing process is summarised in Fig. 4.

**Fig. 4.**
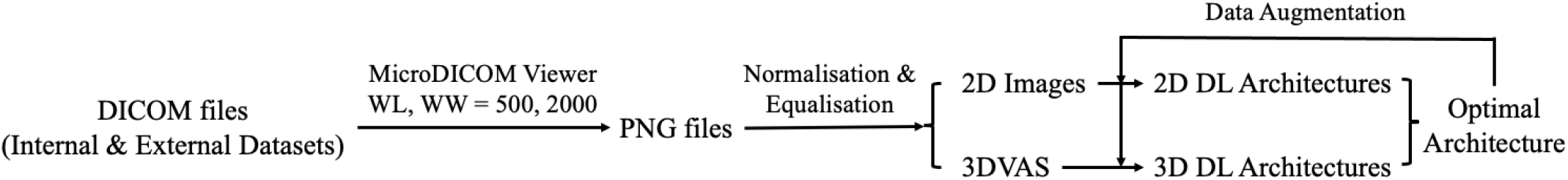
Schematic representation of the data preprocessing pipeline.

### D. Data augmentation

Data Augmentation (DA) techniques have proven to be effective for improving DL models in many different applications [36]. The DA applied in this work consisted of: (i) horizontal and vertical flipping; (ii) 15% scaling down x-axis; (iii) random rotation between *±*180^*°*^; (iv) 20% probability of randomly adding one type of noise, namely: Gaussian (mean=0, std=0.1), Salt & Pepper (salt probability=pepper probability=0.1), Laplace (mean=0, std=0.1), or Poisson noise (determined by pixel values); (v) Gaussian blur with kernel size = (15, 15) and sigma = (2.0, 3.0); (vi) circular cropping to constrain the field of view to the area of the paranasal sinuses; and, (vii) random cropping with squares, circles, and triangles (Fig. 5). The last two techniques were specifically designed to enhance generalisation to the ED. Moreover, to pay attention to different tissue densities [37], the original images were filtered using six window sizes, all different from the default one. Three of them are custom-made: [330, 350], [-30, 30] and [-100, 900]; and the remaining three are the typically predefined skull window ([25, 95]), bone window ([300, 2500]) and mediastinum window ([10, 450]).

**Fig. 5.**
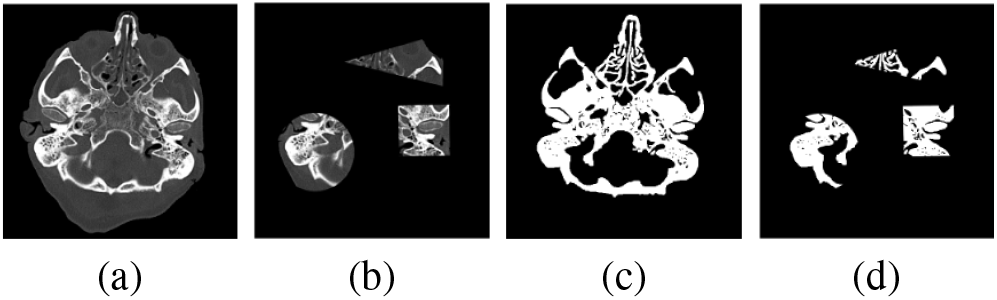
DA: random cropping with squares, circles, and triangles. (a) and (c) are the original slices and their segmented counterparts. (b) and (d) are the corresponding pairs after random cropping.

### E. Evaluation metric

The DICE score was proposed to effectively address data imbalance since anatomical structures or lesions tend to be quite small compared to the background or the rest of the image. The DICE score measures the similarity between two sample images, ranging from 0 to 1, with higher values indicating greater similarity between masks:

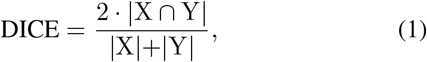

where X and Y represent the pixel values of the two images, respectively. Furthermore, the DICE score is incorporated into the loss function to optimise the different network architectures proposed (see Section II-G). By including the DICE score in the loss function, the model is directly prompted to improve the overlap between the predicted and true masks.

For readability purposes, all DICE scores in this paper were scaled to the range 0-100.

### F. Network architectures

The network architectures used in this work are based on the U-Net [38]. U-Net has consistently demonstrated exceptional performance in medical image segmentation, as evidenced in many studies [23], [24], [27], [28].

A vanilla 2D U-Net architecture was initially used, processing greyscale input images of size 512×512×1. The encoder of the 2D U-Net comprises four blocks, each composed of: (i) a Conv2d with 3×3 kernel; (ii) a ReLU activation; (iii) a dropout layer; (iv) a Conv2D with 3×3 kernel; (v) a ReLU activation; and, (vi) a max pooling of 2×2. The dropout values were set to 0.1 for the first two blocks and 0.2 for the last two. At the bottom of the U-Net architecture, there are two 3×3 convolutional layers and the dropout value was 0.3. The decoder also consists of four blocks, each composed by: (i) a TransposeConv2d with 2×2 kernel and stride of 2×2; (ii) a ReLu activation; (iii) a concatenation of the residual connection; (iv) a Conv2D with 3×3 kernel; (v) a ReLU activation; (vi) a dropout; (vii) a Conv2D with a 3×3 kernel; and, (viii) a ReLu activation final layer. The dropout values were set to 0.2 for the first two blocks and 0.1 for the last two. To generate the greyscale binary masks, the last decoder is composed of two 3×3 convolutional layers also with ReLu activation functions, and are followed by a 1×1 convolution layer with a Sigmoid activation function. The model was optimised using the Adam optimiser with 0.0001 as the learning rate. The batch size was 32, and the number of training epochs was 10. Regarding the vanilla 3D U-Net, same architecture is used using Conv3D instead and 16 batch size and 60 epochs.

In addition, different backbones, such as pre-trained ResNet50 [39] and VGG16 [40], were used as they have achieved excellent results in several studies [41], [42] when applied to medical images segmentation.

Finally, to compare with current large segmentation models, Segment Anything Model (SAM) model [43] released in 2023, alongside the Medical Segment Anything Model (MedSAM) [44] published in 2024 were evaluated. MedSAM is built on the SAM architecture, but specifically adapted for semantic segmentation of medical images. SAM employs a transformer-based architecture, which has proven to be highly effective in a wide range of segmentation tasks. These general-purpose segmentation models have been used as zero-shot predictors.

After the segmentation procedure, the results were post-processed by removing small isolated areas representing non-actual osseous structures and filling in undesired holes, thereby preserving meaningful segmented regions. Post-processing helps eliminate errors that occur during the segmentation process, improving the overall quality of the segmentation results [27].

### G. Loss functions

Two different loss functions were proposed: the Hybrid Loss (HL) and the Assymetric Unified Focal Loss (AUFL).

#### 1) Hybrid Loss

HL combines the DICE loss with the Focal loss function. It is widely used in image segmentation [45], such as in SAM [43].

The DICE loss is derived from the DICE score being ℒ_DICE_ = 1 *−* DICE. To improve the stability of the gradient during training [46], the Focal loss is incorporated. It introduces a modulating factor *γ*, which dynamically scales the cross-entropy loss (ℒ_CE_), reducing the weight of easily distinguishable samples and emphasising the hard-to-distinguish ones. The expression for the ***α***-balanced version of the Focal loss is shown in Eq. (2), where ***α*** represents the class weight vector, **p**_*i*_ is the predicted probabilities vector for each class *p*_*i*,*c*_, where *C* is the number of classes and *i* the pixel pointer [47]. Hence, *p*_*i*,*c*_ is the probability of the *i*-th pixel to belong to class *c*.

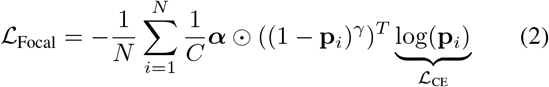

where *N* denotes the number of pixels in an image. Therefore, the HL function is defined as in Eq. (3).

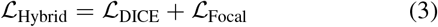

#### 2) Asymmetric Unified Focal Loss

This loss function consists of a modified Focal loss and a modified Focal Tversky loss. According to [46], this loss function resolves the issue of excessive hyperparameters in the original Tversky and Focal loss functions, as well as the convergence problems of the Focal loss at the end of training. Furthermore, through selective enhancement or suppression by focal parameters, asymmetry allows for different losses to be assigned to each class, thereby overcoming the harmful suppression of rare classes and the enhancement of background ones. The expression for the modified Focal Loss (ℒ_*mF*_) is defined in Eq. (4) [46], where the parameter ***α*** in Eq. (2) is replaced with a constant *δ*.

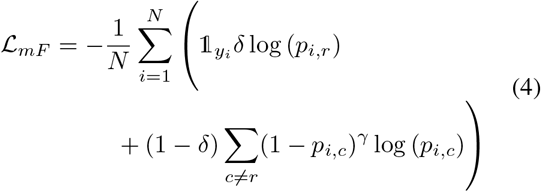

where *y*_*i*_ is the target class for the *i*-th sample (pixel) and 𝟙_*y*_*i* is an indicator function that takes value 1 if *y*_*i*_ = *r* and 0 otherwise.

On the other hand, the modified focal Tversky loss (ℒ_mFT_) can be expressed as in Eq. (5) [46], where *γ <* 1 increases the focus on harder examples, and mTI is the modified Tversky index [46], [48].

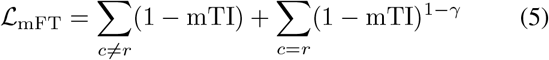

Unlike other loss functions, the Tversky index and its modified version were designed for only two classes: *c* = 1 for the foreground and *c* = 0 for the background. Therefore, mTI is expressed as (Eq. (6)):

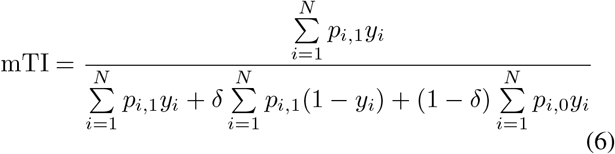

where *y*_*i*_ is the pixel’s target value, which takes 1 for the foreground and 0 for the background.

Using the previous definitions, the AUFL (ℒ_*AUF*_) can be expressed as (Eq. (7)) [46]:

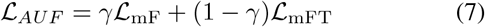

In this work, the hyperparameters of the AUFL were set as: *λ* = 0.5, *δ* = 0.6 and *γ* = 0.5 as proposed by the authors [46].

### H. Experimental protocol

Different experiments were carried out to identify the most suitable scheme for automatic segmentation of the osseous structures of the paranasal sinuses. Three experimental phases were proposed, namely: EP A, EP B, and EP C. They are summarised in Table III, and illustrated graphically in Fig. 6.

**TABLE III.** SUMMARY OF THE EXPERIMENTAL PROTOCOL.

|  | Exp. | Subset | DL Methods | Network architecture | Loss function | DA |
| --- | --- | --- | --- | --- | --- | --- |
| **EP A** | E1 | S1 | Supervised | 2D U-Net (vanilla, ResNet50, VGG16) | HL | No |
|  | E2 | S1 | Supervised | 3D U-Net (vanilla, ResNet50, VGG16) | HL | No |
|  | E3 | S1 | Supervised | Optimal | HL/AUFL | Yes |
| **EP B** | E4 | S1 | Zero-shot | SAM & MedSAM | - | No |
| **EP C** | E5 | S1 & S2 | Supervised | Optimal | Optimal | Yes |
|  | E6 | S1 & S2 | Semi-supervised | Optimal | Optimal | Yes |
|  | E7 | S1 & S2 & S3 | Semi-supervised | Optimal | Optimal | Yes |

**Fig. 6.**
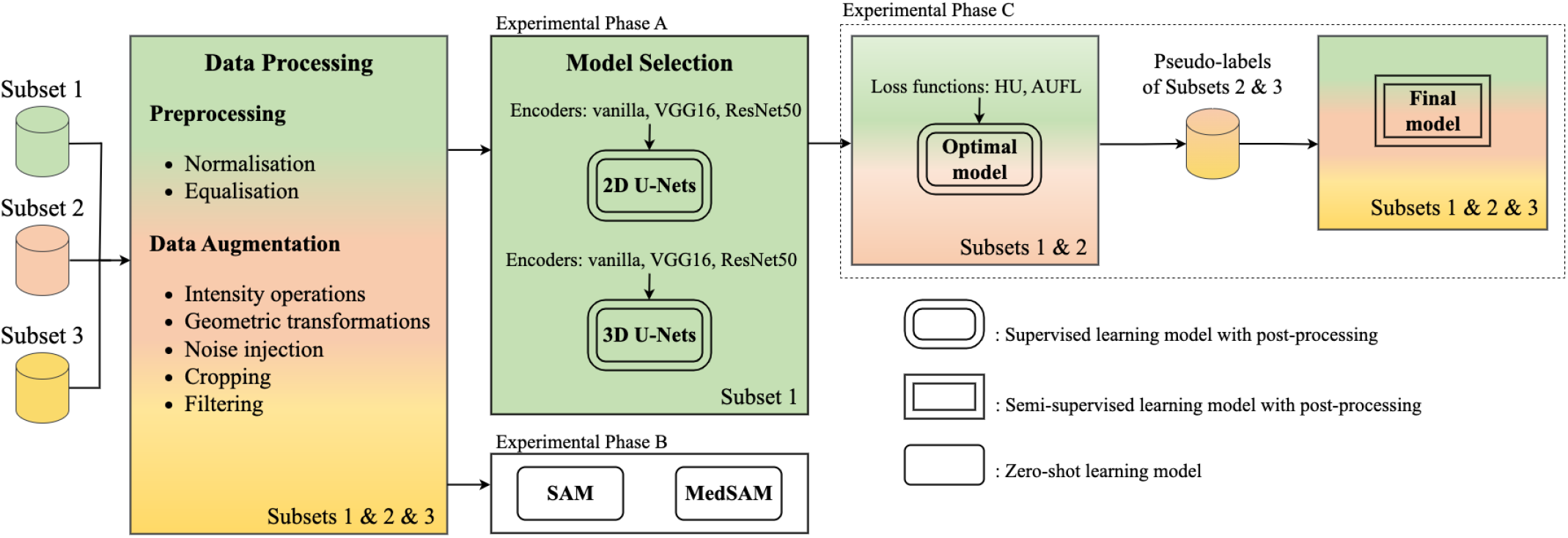
Outline of the overall experimentation protocol. Colours are an indication of the subset used for each EP being S1 green, S2 orange and S3 yellow.

The experiments were carried out using a high-performance computer based on an ASUS^®^ board, with an AMD^®^ EPYC 7313 3.000 GHz CPU, 512 GB of RAM, and four NVIDIA^®^ Quadro RTX A6000 GPUs with 48 GB of VRAM.

1. *EP A:* The goal of this phase was to determine the optimal architecture for the proposed task. Experiments in this phase (E1, E2, and E3) were trained and tested using S1 from the ID (576 slices from 9 patients). A 9-folds Cross-Validation (CV) was conducted with a 7:1:1 distribution for training, validation, and testing. Thus, patients used for training were never used for validation or testing purposes.
  a. *E1:* The E1 used 2D U-Nets. The default vanilla U-Net and two additional backbones (VGG16 and ResNet50) were used. The HL function was used for evaluation.
  b. *E2:* The E2 used 3D U-Nets, also employing the default encoder, VGG16, and ResNet50. The HL function was used for evaluation.
  c. *E3:* Based on the results from E1 and E2, the best network was further improved using the DA techniques and loss functions presented in Section II-G.
2. *EP B*: The goal of this phase was to compare the performance of the best model from EP A with respect to general-purpose pre-trained state-of-the-art segmentation solutions.
  a. *E4*: In E4, the SAM and MedSAM models were tested in the same patients as in EP A, i.e., S1, in a zero-shot procedure.
3. *EP C:* This EP aimed to improve the model’s generalisation to an out-of-distribution dataset. Experiments 5, 6, and 7 (E5, E6 and E7) were conducted using all patients labelled from S1 and different selections from Subsets 2 and 3. To maximise the number of training samples, no validation set was included. All experiments in this phase used the best architecture and loss function identified from EPs A and B. The training time per epoch ranged between 5-16 hours depending on the experiment.
  a. *E5*: In E5, four models were trained using all samples from S1 plus three of the four available from S2. The remaining patient from S2 was used for testing, resulting in a 4-fold CV. In total, 696 slices from 13 patients were used for training and testing purposes.
  b. *E6*: In E6, and based on the study performed in E5, a semi-supervised learning strategy was developed. All unlabelled slices of S2 (221 slices) were pseudo-labelled using the model obtained in E5 for each fold. This semi-supervised approach expanded the available data to 917 slices from 13 patients. The strategy is similar to that followed in [49], [50].
  c. *E7:* In E7, 30 slices manually selected from each patient of S3 were pseudo-labelled with the E6 model obtaining 810 pseudo-masks for training. Then, following same CV as in E5 and E6, a final model was trained. To balance the number of labelled and pseudo-labelled samples while minimising error propagation, the training included 12 labelled patients and 12 pseudo-labelled. Each epoch randomly selected 12 pseudo-labelled patients from S3, thus increasing the variability. Furthermore, to decrease computational burden, a transfer learning strategy was used, starting from the models obtained in E6 for each fold.

## III. RESULTS

The results obtained for all experiments carried out in all phases (EP A, EP B and EP C) are presented below.

### A. Experimental Phase A

Table IV illustrates the results obtained for E1 and E2. According to the results, the use of 2D and 3D U-Net architectures has a negligible impact on the results, as the DICE coefficients are very similar for both architectures. However, in terms of training time, 2D architectures converged faster than 3D ones (0.15 min. per epoch for 2D models, and 0.3 min. for 3D ones).

**TABLE IV.** DICE SCORES OBTAINED FOR E1 AND E2.

| Exp. | U-Net | Vanilla * | ResNet50 * | VGG16 * |
| --- | --- | --- | --- | --- |
| E1 | 2D | 90.20 ± 2.3 | 91.55 ± 1.6 | 91.70 ± 1.6 |
| E2 | 3D | 90.71 ± 1.3 | 89.88 ± 1.3 | 90.77 ± 1.5 |

\* mean±std

In any case, among the six architectures tested, the 2D U-Net with VGG16 encoder provided better DICE scores (absolute improvement of 0.93). Consequently, this architecture was chosen for E3.

Table V illustrates the results obtained for E3. The DICE score obtained using an AUFL loss function was, on average, 0.05 higher than that achieved using HL. On the other hand, DA has not reported significant improvements. DA increased the number of training samples in S1 from 576 to 21,888 slices (i.e., each slice derived in 38 variants). As a consequence, the training time for each epoch raised up to 90 min. Despite DA having not significantly improved the model, it was retained for the next experiments to ensure better generalisation.

**TABLE V.** DICE SCORES OBTAINED FOR E3.

| Exp. | DA | HL loss * | AUFL loss * |
| --- | --- | --- | --- |
| E3 | Yes | 91.24 ± 1.6 | 91.31 ± 1.4 |
| E3 | No | 91.70 ± 1.6 | 91.73 ± 1.7 |

\* mean±std

In view of these results, the model selected for EP C (i.e., validation with the ED) used: (i) 2D U-Net with a VGG16 encoder; (ii) an AUFL loss function; and, (iii) DA techniques.

### B. Experimental Phase B

The qualitative results obtained using SAM and MedSAM are shown in Fig. 7, where each colour represents a different class. The SAM model (Fig. 7.b) was able to segment the background and the external contour of the patient’s head. However, it struggled to detect the osseous structures. For example, only part of the frontal sinus was detected, while the other paranasal sinuses were not identified.

**Fig. 7.**
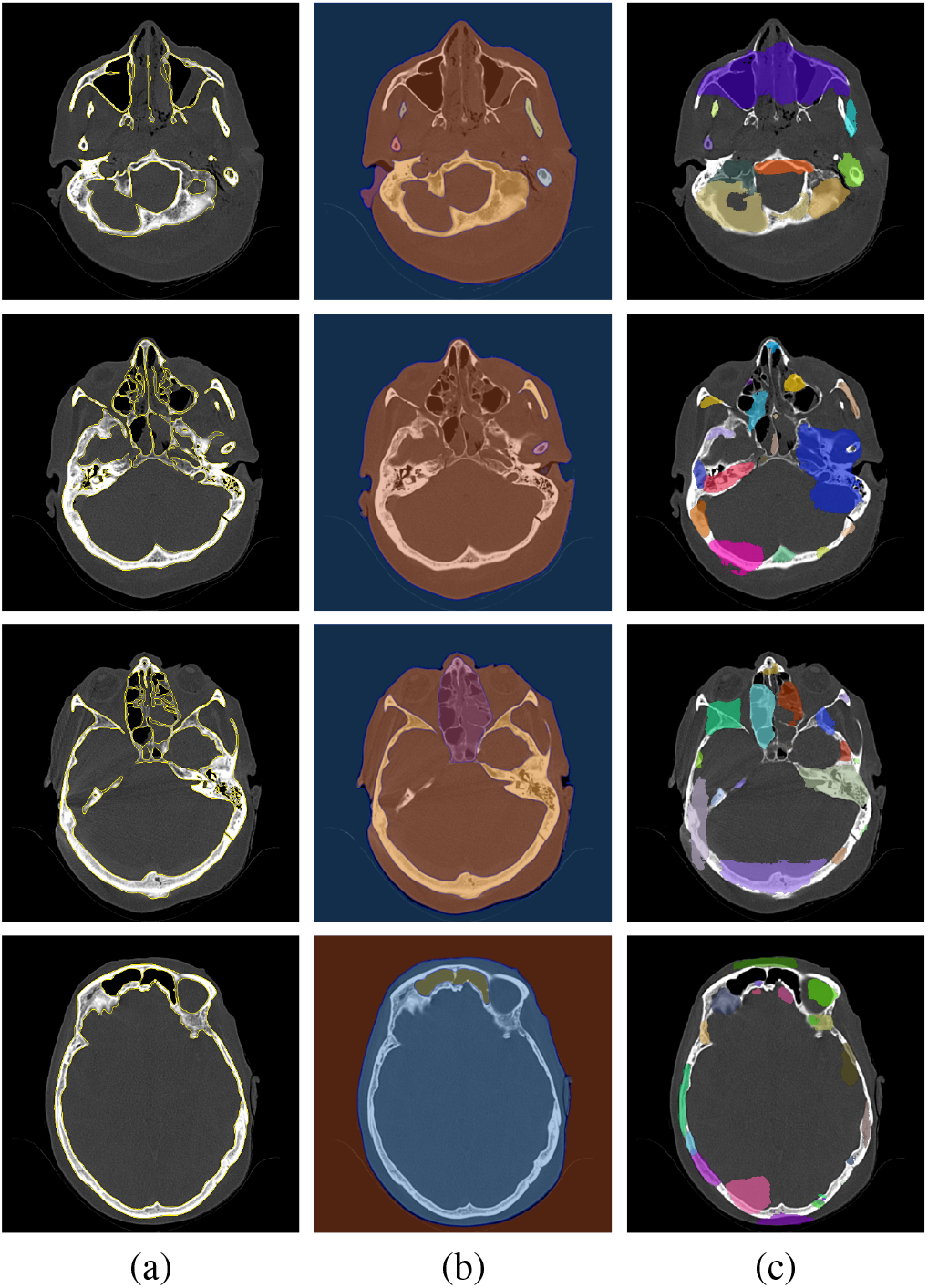
Examples of masks generated by the SAM and MedSAM models. (a) Original CT slices with manual contour lines (ground truth). (b) Results provided by SAM. (c) Results obtained by MedSAM.

**Fig. 8.**
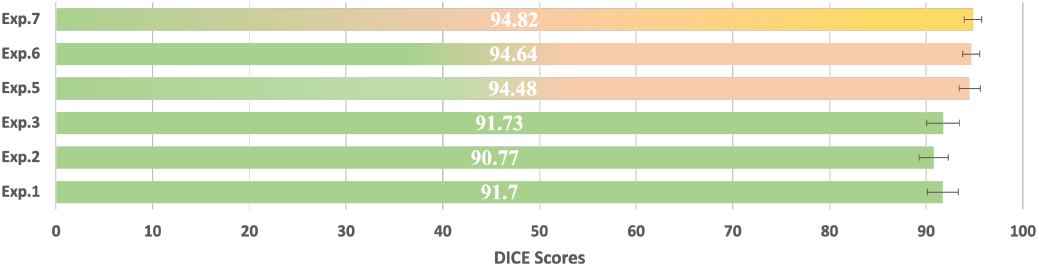
Summary of the best results for each experiment. The colors of the bars represent the different subsets used in the training set (green: S1, orange: S2, yellow: S3).

A similar pattern was observed with the MedSAM model (Fig. 7.c). Despite the visual prompts providing to MedSAM, the segmentations were not accurately aligned with the osseous structures.

### C. Experimental Phase C

In this phase, the generalisability of the model was evaluated. DICE scores reported in this section were calculated using S2.

Table VI presents the results obtained for E5, E6 and E7. The results obtained show that the models developed in E3 were able to generalise to the external S2: the DICE scores even improved with respect to those in E1-E3. Fig. 9 graphically presents several examples of predictions from E5, E6 and E7.

**TABLE VI.**
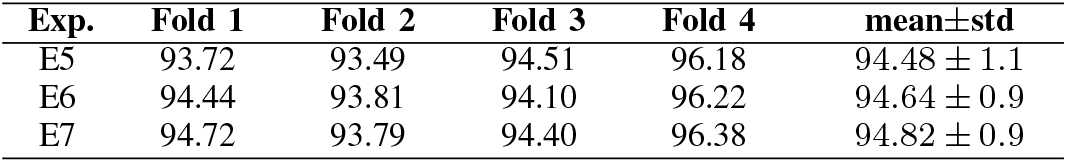
DICE SCORES OBTAINED FOR E5, E6 AND E7.

| Exp. | Fold 1 | Fold 2 | Fold 3 | Fold 4 | mean±std |
| --- | --- | --- | --- | --- | --- |
| E5 | 93.72 | 93.49 | 94.51 | 96.18 | 94.48 ± 1.1 |
| E6 | 94.44 | 93.81 | 94.10 | 96.22 | 94.64 ± 0.9 |
| E7 | 94.72 | 93.79 | 94.40 | 96.38 | 94.82 ± 0.9 |

**Fig. 9.**
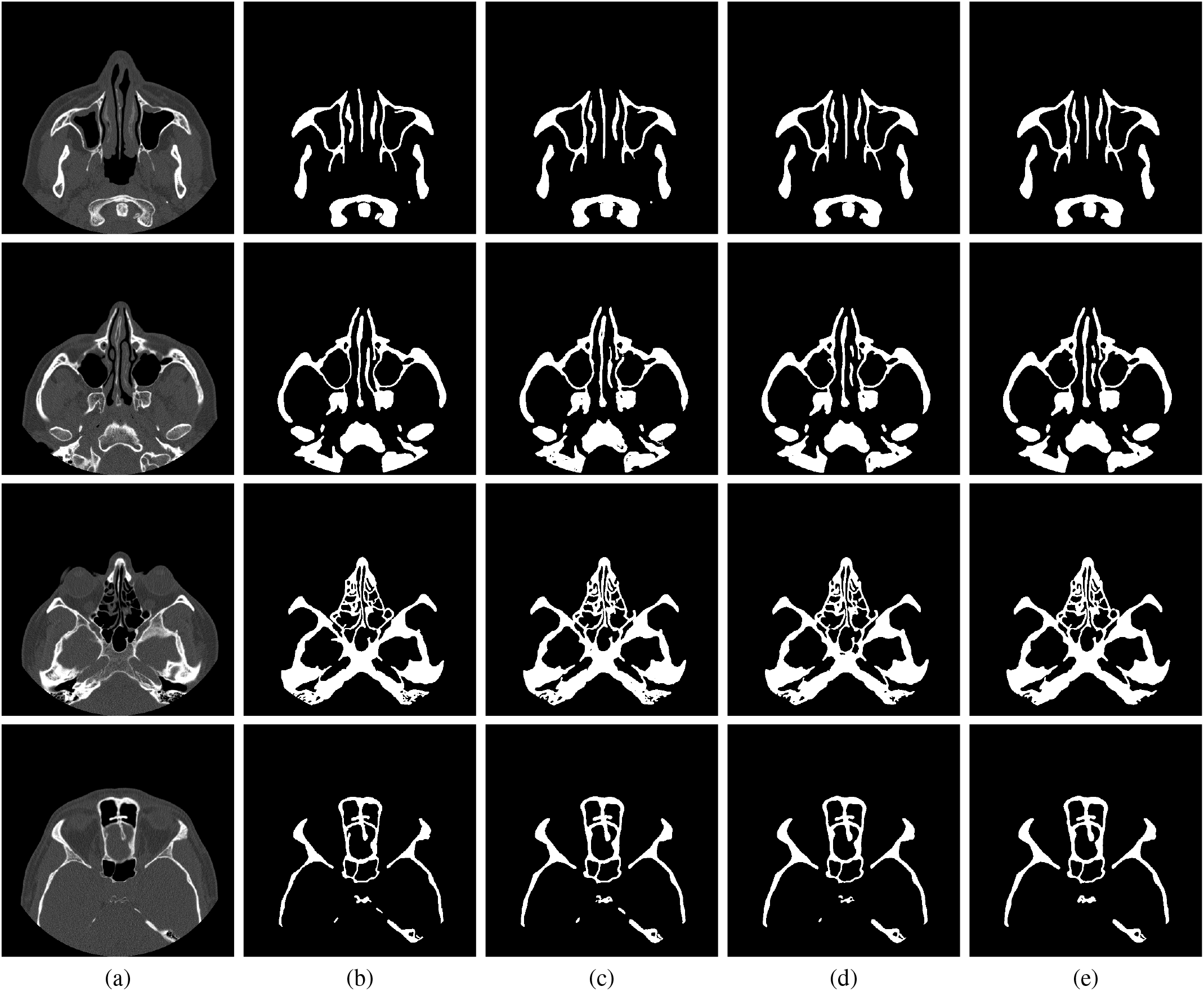
Results from E5 and E6. (a) Original CT slices. (b) Ground truth. (c) & (d) & (e) Masks obtained in E5, E6, and E7, respectively.

Furthermore, with respect to E5, results in E6 and E7 showed a modest increase in the mean DICE and a reduction in its standard deviation. This suggests that the semi-supervised and transfer learning strategies followed in these experiments were able to improve the generalisability of the model.

The impact of the post-processing stage in the E7 results is illustrated in Fig. 10. Despite visible minor errors, post-processing effectively improved the reliability of the system and thus is considered a crucial step.

**Fig. 10.**
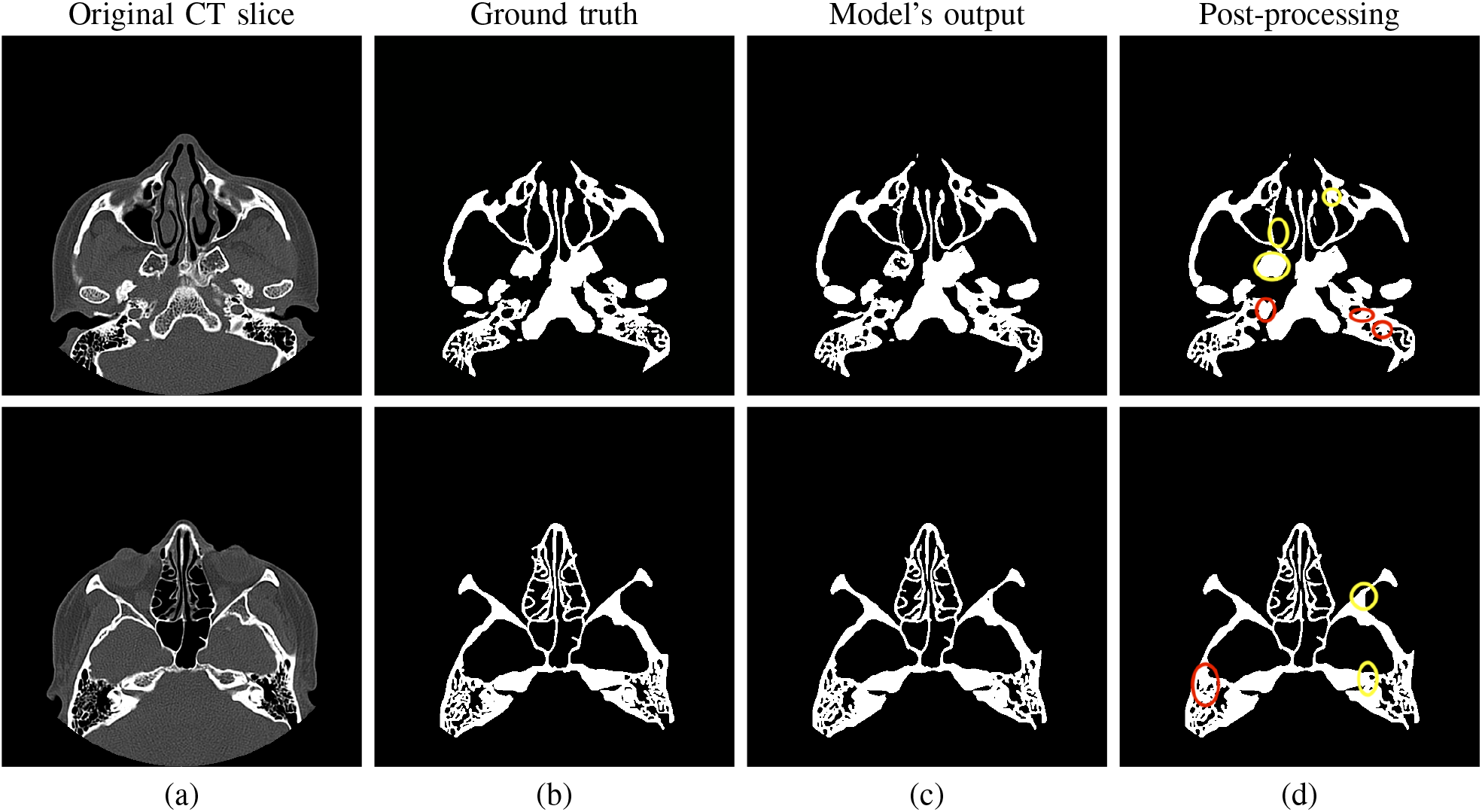
Illustration of results from E7. (a) Original CT slice. (b) Ground truth. (c) Results without post-processing. (d) Results post-processed (yellow circles indicate areas where post-processing successfully addressed false positives or true negatives; red circles indicate errors).

Finally, Fig. 11 illustrates different errors committed by the system trained in E7. Comparison of the ground truth with the predicted masks reveals that, although they were very similar for large structures, the predicted masks exhibit certain errors in fine detail. This discrepancy was particularly pronounced along the edges of the osseous structures around the ethmoid sinuses, and in other small anatomical structures.

**Fig. 11.**
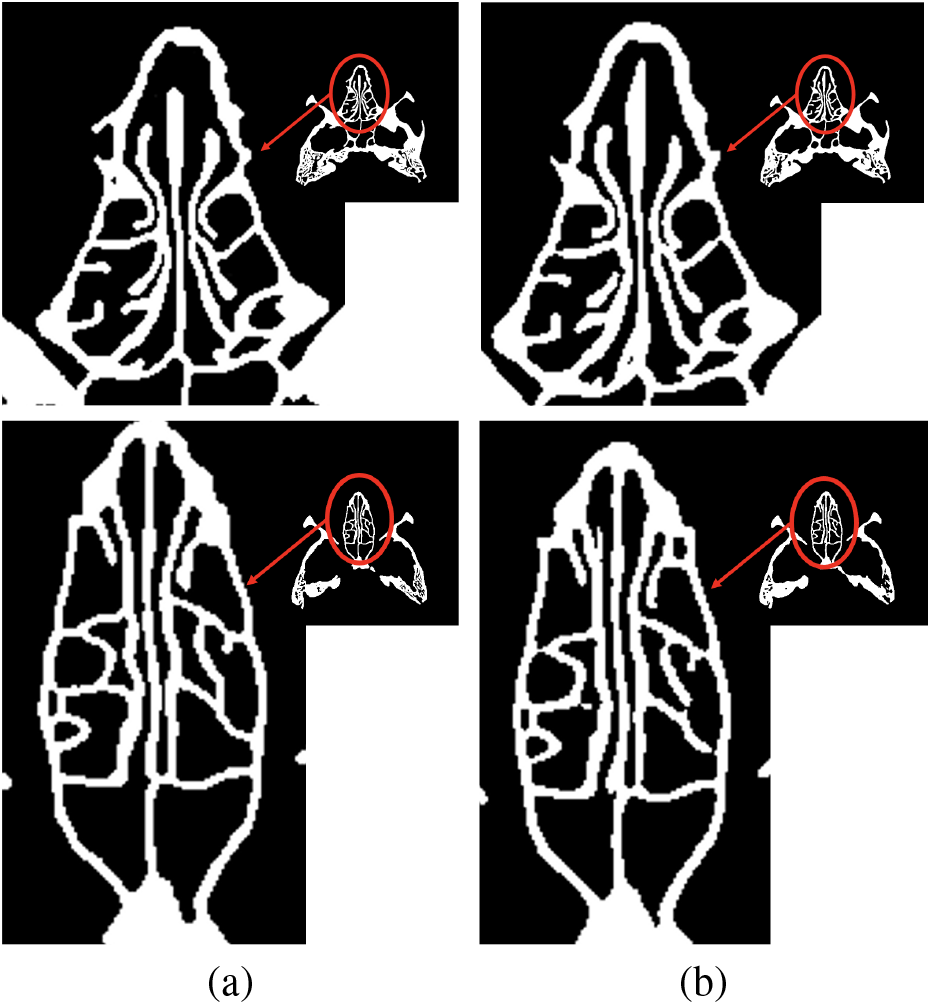
Zoomed-in view of typical errors in the segmentation process. (a) Ground truth. (b) Results from E7.

Regarding training time, E5 required 150 min. per epoch. This is attributed to a larger training set —from 576 to 696 slices, resulting in 26,448 slices after DA—. In E6, training took 180 min. per epoch due to a larger training set —917 slices, 34,846 after DA—. In E7, 360 slices from S3 were randomly added to the training set for each epoch, increasing the number of slices —to 1,277, 48,526 after DA—. Consequently, the training time per epoch increased to 270 min.

For the sake of comparison, the results for all experiments in terms of DICE are graphically summarised in Fig. 8.

## IV. DISCUSSION

Automated segmentation of the OSPS can enhance a computer-assisted workflow for preoperative virtual surgical planning, CAD/CAM-assisted or neuronavigated surgery, and postoperative verification of the outcomes. While accurate segmentation of the osseous structures is often the starting point for virtual planning, it provides valuable information for computer-assisted planning and surgery. For example, in combination with neuronavigators, can offer crucial intraoperative information to the surgeon by providing visual and/or haptic feedback during surgery, or can provide extra information for the creation and precise placement of patient-specific implants (e.g., in reconstructing complex orbital defects). Additionally, in septoplasty, automated segmentation of the osseous structures can aid in virtual surgery planning and execution.

Besides, the rapid advancements in augmented and virtual reality applications are expected to further promote computer-assisted craniomaxillofacial surgery. To fully exploit these developments, it is crucial to have a detailed and precise segmentation of the OSPS is crucial.

However, to our knowledge, no studies have specifically targeted personalised segmentation of the OSPS. Instead, existing research has concentrated mainly on segmenting the upper airways within the paranasal sinuses [17], leaving a notable gap in the precise delineation of the osseous structures, which is considered a much more challenging task.

In addition, the absence of open datasets significantly limits not only the research about specific segmentation techniques for delineating these structures, but also the development of models specifically tailored to these anatomically complex regions. To our knowledge, the corpus used in this study is the first developed for this purpose that has been manually annotated and made openly available.

To this respect, and to support diverse research needs, we have developed —and made available— four different models: (i) trained on nine ex vivo patients; (ii) trained on all patients with manual segmentations (from both ID and ED); (iii) semi-supervised model incorporating manual and pseudo-labels from 13 patients; and, (iv) a model trained on 13 manually segmented patients plus 27 pseudo-labelled.

In EP A with S1, the best DICE coefficient was 91.73 *±* 1.7. These results were obtained using a 2D U-Net with a VGG16 encoder. However, the difference with respect to the other U-Net architectures tested is scarce, as also reported in [24] for a different application domain.

In terms of comparison, the results obtained are lower than those reported in [23], where authors reached DICE values of 96.0 for the upper skull. However, the comparison is not straight due to differences in the target volume segmented.

The volume reconstructed in [23] covers the whole upper skull, including structures that are much easier to identify and segment, significantly biasing the DICE. The work in [24] could also be used for comparison purposes. In this work, authors obtained DICE scores of 96.5 for the maxilla and 98.4 for the mandible. However, as in the previous case, the problem posed in this work is less challenging because of the presence of large bones that are much easier to segment. Another potential comparison could be established with the results reported in [27], where the authors obtained a DICE score of 78.8 for the craneal base. The comparison, once again, is not straight due to significant differences in the target structures.

Regarding generalization with a different dataset, the models developed have demonstrated robust capabilities with the ED in S2 and S3. These results contrast with those obtained in [24] (also using an external dataset), where the authors report a substantial decrease of 10 absolute points in the DICE score. This is attributed to the ambitious DA strategy followed in this work, which did not significantly improve scores in EP A, but has enhanced the generalisation capabilities of the model. Furthermore, the semi-supervised learning strategy followed addressed the challenge of insufficient data available, yielding an average increase of 0.34 for the DICE score.

In summary, the highest DICE coefficients achieved between the predicted mask and the ground truth were 91.73*±* 1.7 tested on S1, and 94.82 *±* 0.9 tested on S2. These results are graphically illustrated in Fig. 12, which presents two 3D renderings of the OSPS derived with E3. Fig.12.Top graphically exemplifies the errors committed by the model.

**Fig. 12.**
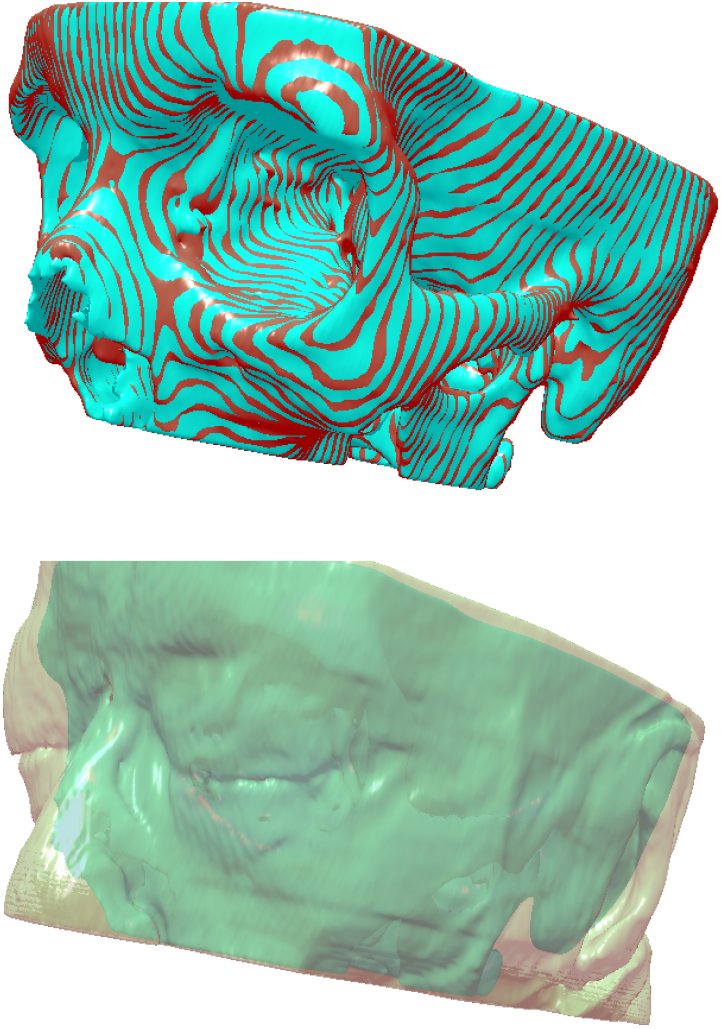
Example of a 3D reconstruction using the models developed in E3. Top: Ground truth in red, and automatic delineation in blue. Bottom: Automatic delineation with the contour of the skin overlayed.

Segmentation modelling of the OSPS still has great potential for improvement. The corpus used is limited, so increasing the sample size with more manually annotated masks could potentially enhance the performance. In any case, new training data could also be generated using a pseudo-labelling strategy similar to that followed in this work.

Additionally, exploring different variants of the U-Net architecture may yield beneficial results. To this respect, U-Net Transformer (UNETR) (based on a Vision Transformer (ViT)) has reported relevant results in other application domains. U-Mamba and U-Kolmogorov-Arnold Networks (U-KAN) are also novel and relevant alternatives [51].

Furthermore, although the current SAM and MedSAM did not perform well for the aimed segmentation task, they remain promising for future research, since they could be integrated as fine-tuning methods after an initial segmentation using the methods proposed in this paper. On the other hand, MedSAM could be used for a transfer learning-based training strategy.

It is important to note that, in this study, segmentation and validation were performed on healthy CT scans (even when the ID corresponds to ex vivo heads). Therefore, altered anatomical structures, such as those resulting from trauma or tumours, could affect the segmentation accuracy of the models.

Another potential limitation of the current study is the need for an in-depth study of possible variability factors in the training data, such as the scanner model or manufacturer and the patient’s age, sex, or race.

Finally, it should be mentioned that the manual segmentation of the dataset was reviewed by a single expert, thus models are expected to reproduce one single segmentation criterion. Including more experts would lead to more variability, and might lead to more robust models (although not necessary with a better DICE).

## V. CONCLUSIONS

Six different models based on the U-Net architecture were developed for the segmentation of the OSPS from axial CT scans. They are based on the 2D and 3D U-Net architectures and their corresponding variants using VGG16 and ResNet50 encoders. Although vanilla U-Net has the advantage of a simple structure and less training time, its variants using VGG16 and ResNet50 provided better results, taking advantage of their pre-training. The best results were obtained using the 2D U-Net with a VGG16 backbone.

The impact of DA techniques and potential improvements due to loss functions were also evaluated. The results show that the wide range of applied DA techniques significantly improve the models’ generalisation. On the other hand, AUFL provided slightly better results for the problem posed.

The impact of a pseudo-labelling strategy on the generalisation ability of the models was also evaluated. The results obtained show that this technique can improve the accuracy of the developed models.

Models were trained using a relatively small dataset of 13 patients. Despite this limitation and the complexity of the task, the results suggest that the models developed perform well even with small sample sizes. Improving the model by scaling to a wider dataset is straight, but would require more expensive computational resources, since the best model of those trained required almost two weeks of uninterrupted computing.

Additionally, the SAM and MedSAM models, recently released for general-purpose segmentation problems, exhibited poor performance. The limited ability of these models to address this task is attributed to SAM’s limitations in recognising fuzzy boundaries and, to the lack of osseous structures around the paranasal sinuses region in MedSAM’s training data [44]. To address this, we have made all models trained in this work freely available, allowing them to be used as transfer learning backbones for other bone segmentation applications.

## ACKNOWLEDGMENTS

The authors thank C. Hoyos-Barceló, G. Pérez-de-Arenaza-Pozo and J. C. Puerta-Acevedo for collaborating in the manual annotation of the images. The authors gratefully acknowledge the Universidad Politécnica de Madrid for providing computing resources on the Magerit Supercomputer.

## Footnotes

1 Sinus scans are recommended in case of sinus cancer and other malignant and metastatic tumours, inflammatory diseases of the sinuses, trauma, and preoperative assessments.

